# Impact of early childhood malnutrition on the adult brain function: an ERP study

**DOI:** 10.1101/782698

**Authors:** Kassandra Roger, Phetsamone Vannasing, Julie Tremblay, Maria L. Bringas Vega, Cyralene P. Bryce, Arielle G. Rabinowitz, Pedro A. Valdés-Sosa, Janina R. Galler, Anne Gallagher

## Abstract

According to the World Health Organization, 45% of deaths among children under five years of age are caused by malnutrition, which impacts more than 224 million children globally. The Barbados Nutrition Study (BNS) is a 50+ year longitudinal study on a Barbadian cohort with histories of moderate to severe protein-energy malnutrition (PEM) limited to the first year of life and a healthy comparison group. We have previously used quantitative electroencephalography (EEG) to highlight differences in brain function during *childhood* (lower alpha1 activity and higher theta, alpha2 and beta activity) between participants who suffered from early PEM and controls. In order to determine whether similar differences between the PEM and control groups persisted into *adulthood*, our current study used recordings obtained during a Go-No-Go task in a subsample of the original BNS cohort (N=53) at ages 45-51 years. We found that previously malnourished adults (*n*=24) had a higher rate of omission errors on the task relative to controls (*n*=29). Evoked-Related Potentials (ERP) were significantly different in participants with histories of early PEM, who presented with lower N2 amplitudes (p<0.05). These findings are typically associated with impaired conflict monitoring and/or attention deficits and may therefore be linked to the attentional and executive function deficits that have been previously reported in this cohort in childhood and again in middle-adulthood.

**Highlights:**

- Childhood malnutrition increases risk of brain function alterations.
- There is a need to investigate the evolution of those outcomes later in life.
- Adults who suffered childhood malnutrition undertook a Go-No-Go task during EEG.
- Task performance and N2 amplitude were reduced in malnutrition group (vs control).
- First evidence of adult brain function alteration following childhood malnutrition.

## 1. Introduction

More than two-hundred million children under age five are affected by malnutrition worldwide, which makes this condition a critical global health concern (UNICEF, 2019). Protein-energy malnutrition (PEM) is a specific type of malnutrition defined as an acute caloric deficit due to deficiency of all macronutrients, and micronutrients in some cases (Atassi, 2019; Grover and Ee, 2009; Morley, 2018). Determining the effects of this important health problem early in life is therefore a priority.

While malnutrition has deleterious effects on cognitive abilities (Berkman et al., 2002; Champakam et al., 1968; Galler et al., 1984a, 1983b; Kar et al., 2008; Mendez and Adair, 1999; Upadhyay et al., 1989a), it has also been associated with various effects on motor skills, social abilities and mental health (Galler et al., 1983a, 1984b, 1985; Upadhyay et al., 1989b), with many of these effects persisting into adolescence and even adulthood (Galler et al., 2012a, 2012c; Hoorweg and Stanfield, 1976; Waber et al., 2014b, 2014a; Walker et al., 2007). Using electroencephalography (EEG), brain function alterations have also been linked to early childhood malnutrition (for a review see Gladstone et al., 2014). Although these findings were replicated in several studies (Baraitser and Evans, 1969; Griesel et al., 1990; Stoch et al., 1963), few of them assessed the children at later timepoints in childhood. Bartel and colleagues (1979) studied malnourished children five to ten years after their hospitalization and confirmed that their rest EEGs still exhibited significantly diminished alpha and increased theta and delta frequencies compared to a control group. It is clear from these behavioral and EEG findings that malnutrition at a young age has important acute deleterious effects on early brain function, cognition and behavior. However, the effects of malnutrition on brain function in adulthood are still unknown.

The Barbados Nutrition Study (BNS) is a 50-year longitudinal study that has followed a Barbadian cohort hospitalized during the first year of life for a single episode of moderate to severe protein-energy malnutrition (PEM; (Galler et al., 1983a, 1987) based on the Gomez classification (Gomez et al., 1955), and matched healthy controls who were former classmates of the PEM participants. The objectives of this study are to characterize the medical, neuropsychological, behavioral and brain effects of early PEM over the lifespan. The cohort comprises participants that were originally recruited between 1967 and 1972 when they were hospitalized as infants and subsequently enrolled in a government program that provided health and nutrition monitoring until the children reached 12 years of age. The malnutrition episode was restricted to the first year of life and all children in the PEM group achieved complete catch-up in physical growth by adolescence (Galler et al., 1987). Control participants were classmates of the PEM children and were matched for age, sex and handedness. Neuropsychological and psychiatric assessments revealed many cognitive, behavioral and mental health impairments associated with early childhood PEM, including lower IQ, conduct problems and higher prevalence of affective and depressive symptoms, as well as attention deficits (Galler et al., 2012b, 2010, 1983a, 1983b). Additionally, most participants in the PEM group exhibit persistent attention and executive problems during childhood, adolescence and adulthood compared to the control group (Galler et al., 2012c, 1983a; Galler and Ramsey, 1989). In a recent study, we reported on brain function alterations in the resting state activity of previously malnourished 5-11year-old children from the BNS cohort: specifically, an excess of theta, alpha2 and beta frequencies, and a decrease of alpha1, all suggesting a maturational lag in cortical development despite complete nutritional rehabilitation and demonstrated catch-up growth (Taboada-Crispi et al., 2018). However, little is known about the long-term effects of early childhood malnutrition on brain function in adults from this cohort.

The present study arises from an international Canada-United States-Cuba-Barbados collaboration. Our aim was to study the persistence of brain function alterations in adults who experienced moderate to severe PEM during the first year of life by comparing the brain activity in adults from the PEM and the control groups of the BNS cohort using Evoked-Related Potentials (ERPs). Considering the attention and executive impairments previously reported in the PEM group (Galler et al., 2012c, 1983a; Galler and Ramsey, 1989), we administered a Go-No-Go attention task during EEG recordings to specifically isolate those altered processes. In a Go-No-Go task, two ERPs components are typically studied, namely the N2 and P3 components. N2 is defined as the largest fronto-central negative polarity deflection between 200 and 450 ms, and P3 is defined as the largest fronto-central positive polarity deflection between 350 and 600 ms (Grane et al., 2016; McLoughlin et al., 2010; Schmüser et al., 2016; Woltering et al., 2013). N2 and P3 are respectively associated with conflict monitoring (i.e. conflict between the prepotent response and required response) and inhibition response (i.e. cancellation of a planned or prepotent response) processes in an attention/inhibition control task such as Go-No-Go, Stop signal, Stroop or Flanker task (Braver et al., 2001; Smith et al., 2008). Those components have been widely studied and are thought to be altered in populations with attention and executive impairments like ADHD (Grane et al., 2016; Johnstone et al., 2013; McLoughlin et al., 2010; Schmüser et al., 2016; Woltering et al., 2013) and schizophrenia (Araki et al., 2016; Doege et al., 2010). In the current study, we hypothesised that the PEM group would show smaller N2 and P3 component amplitudes compared to the healthy control group.

## 2. Materials and Methods

### 2.1. Site of Study

All study participants were born in Barbados, an English-speaking Caribbean country, between 1967 and 1972. Barbados has a population of 285,000 persons, primarily of African origin (92%), and the majority are lower middle class. The literacy rate is 99% and school attendance is obligatory to 17 years. In 1970, during the period when the sample was born, the infant mortality rate in Barbados was 46 per 1000 live births, compared with 7.8 per 1000 live births today (UNICEF, 2018).

### 2.2. Participants

Fifty-five adults, all born between 1967 and 1972, were recruited from the Barbados Nutrition Study (BNS) cohort during June-July 2018 (Galler et al., 1983b, 1983a; Ramsey, 1980) and were approximately 45-51 years when tested. Briefly, the PEM group participants were hospitalized with moderate to severe PEM (Gomez et al., 1955) limited to the first year of life. Inclusion criteria were: (1) birth weight >2500 g (2) no pre- or postnatal complications (3) Apgar scores >8, (4) no encephalopathic events during childhood and (5) no further malnutrition after age one. The children were enrolled in an intervention program until 12 years of age, that provided subsidized foods, nutrition education, home visits, medical care and a preschool program two to three mornings a week (Ramsey, 1980). Healthy controls came from the same classrooms and met the same inclusion criteria but had no histories of malnutrition. They were matched to the PEM group by age, gender and handedness (to allow comparison of neurodevelopmental measures). In the current study, two participants from the PEM group had to be excluded, one due to heavy alcohol consumption the night before testing, and the other because he could not complete the task. The final sample included 24 PEM participants and 29 controls. Analysis to determine the representativeness of the current sample relative to the original Wave 1 cohort yielded no significant differences between current study participants (*n*=53) and non-participants (*n*=205) who were studied in earlier waves with respect to gender (percent male: *χ*2= 1.93, *p*= 0.17), history of childhood malnutrition (*χ*2= 0.59, *p*= 0.44), childhood ecology (*t*= 1.11, *p*= 0.27), and childhood IQ (*t*= 0.69, *p*= 0.49).

### 2.3. Procedure

EEG recordings took place at the Barbados Nutrition Study centre, Bridgetown, Barbados. The room was air conditioned and the temperature was maintained at 24°C. Participants were seated in a comfortable chair and fitted with a 21-EEG Ag/AgCl electrode active cap, one vertical and one horizontal electro-oculogram (EOG), and one electrocardiogram (ECG). The EEG signal was recorded on the scalp using the actiCHamps amplifier and Brain Vision Recorder Software (Version 1.20, Brain Products, GMbH, Gilching, Germany). EEG channels were positioned according to the 10-20 system (Fz, Cz, Pz, Oz, Fp1/2, F3/4, C3/4, P3/4, O1/2, F7/8, T3/4, T5/6, plus ground and reference). Impedance was kept under 10kΩ. A near-infrared spectroscopy (NIRS) recording was performed simultaneously with the EEG. However, the NIRS data are not included in the current article and will be reported in a subsequent publication. A Go-No-Go task was performed during the EEG-NIRS recording.

### 2.4. Go-No-Go task

The Go-No-Go task was presented using Presentation (version 20.2, Neurobehavioral Systems; see Figure 1). Participants were instructed to click on a computer mouse as fast as possible using the index finger of their dominant hand whenever a letter appeared on the screen (Go trial), but to withhold from clicking if the letter was an X (No-Go trial). Each letter was presented pseudorandomly for 500 ms and disappeared as soon as a motor response was made. The inter-stimulus interval varied randomly between 700 and 1000 ms. The task utilized a block design in which each block of 20 stimuli was interleaved with a resting period that varied randomly between 15 and 21 sec. A block design was used rather than a single trial paradigm to accommodate both the EEG and NIRS signals. The task consisted of 14 No-Go blocks, and 14 Go blocks. The No-Go blocks included 30% of No-Go trials (6 X) and 70% of Go trials (14 letters). The Go blocks included Go trials only. Each participant went through a practice trial of one No-Go block that included feedback before the actual task. Overall, 28 blocks were presented, for a total of 476 Go and 84 No-Go trials (total of 560 letters). Additional blocks were presented so that at least 70 No-Go trials were correctly inhibited to ensure a good signal to noise ratio. Three behavioral variables were computed based on the 28 blocks of the task (without additional blocks): reaction time (RT), No-Go accuracy and Go accuracy.

**Figure 1.**
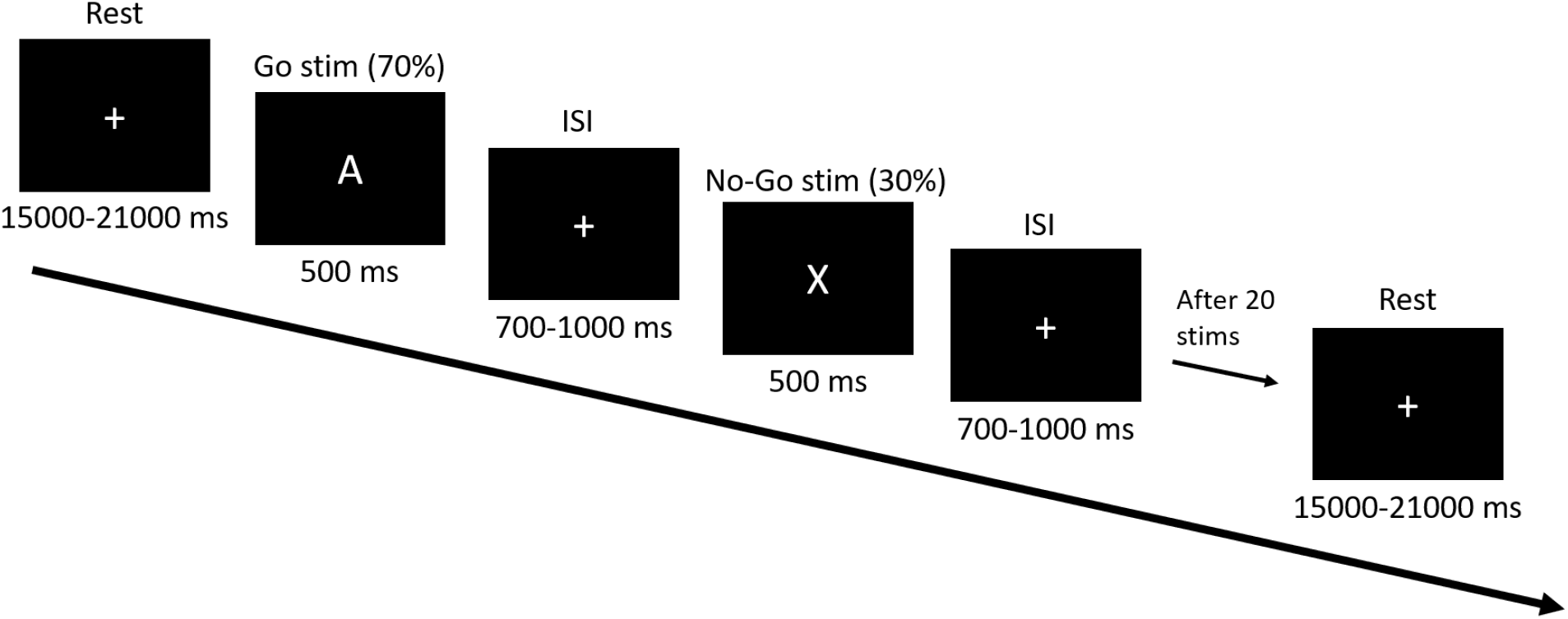
Go-No-Go task design

### EEG recording and data processing

The EEG signal was recorded from the scalp using a 500Hz sampling rate and referenced to electrode FCz. The data pre-processing was done using Brain Vision Analyzer (Brain Products, Munich, Germany). Data were first filtered offline between 0.5 and 35 Hz using a Butterworth filter with a notch filter at 50 Hz to remove any electrical interference. Ocular artefacts were then removed by subtracting the corresponding components using Independent Component Analysis (ICA). The data were then re-sampled at 512 Hz and re-referenced to the average reference. Each trial was segmented from −500 to 1000 ms after stimulus onset. DC detrend was applied to the data to remove signal drift.

Artifact correction was then performed to reject any segment with artifacts for each channel individually. Any segment with a voltage step > 50 μv/ms was removed. The maximum amplitude allowed was 100 μv and the minimum amplitude allowed was −100 μv. A trained Ph.D. candidate (K.R.) then performed a visual inspection of the signal to detect any remaining artifacts and validate the artifact correction. Every incorrect trial, namely every No-Go followed by a motor response or Go without a motor response, was also rejected. A baseline correction was further applied using 200 ms before stimulus onset. Trials were then averaged in Correct No-Go (*M* = 69.6 trials, *SD* = 10.5) and Correct Go trials (*M* = 471.3, *SD* = 67.3).

### 2.6. ERP analysis

For each participant, a semi-automatic peak detection for the N2 and P3 ERP components was first performed on Correct No-Go and Correct Go trials individually on Fz, Cz and FCz electrodes, where the components have previously been reported (Donkers and van Boxtel, 2004; Falkenstein et al., 1999). N2 was defined as the largest fronto-central negative polarity deflection between 200 and 450 ms and P3 was defined as the largest fronto-central positive polarity deflection between 350 and 600 ms (Grane et al., 2016; McLoughlin et al., 2010; Schmüser et al., 2016; Woltering et al., 2013). The peaks were reviewed by a trained Ph.D. student (K.R.) and an expert in EEG processing (P.V.). The mean amplitude ±10 ms around the peak and the latency of each peak was extracted and used for statistical analyses. A topographic T-test (without correction for multiple comparisons) was performed to compare amplitude topographies between Nutrition groups (PEM vs Control). Brain Vision assisted t-tests were computed on amplitude topographies during N2 and P3 components for No-Go and Go conditions individually when group differences were maximal (see Figures 2 and 3).

**Figure 2.**
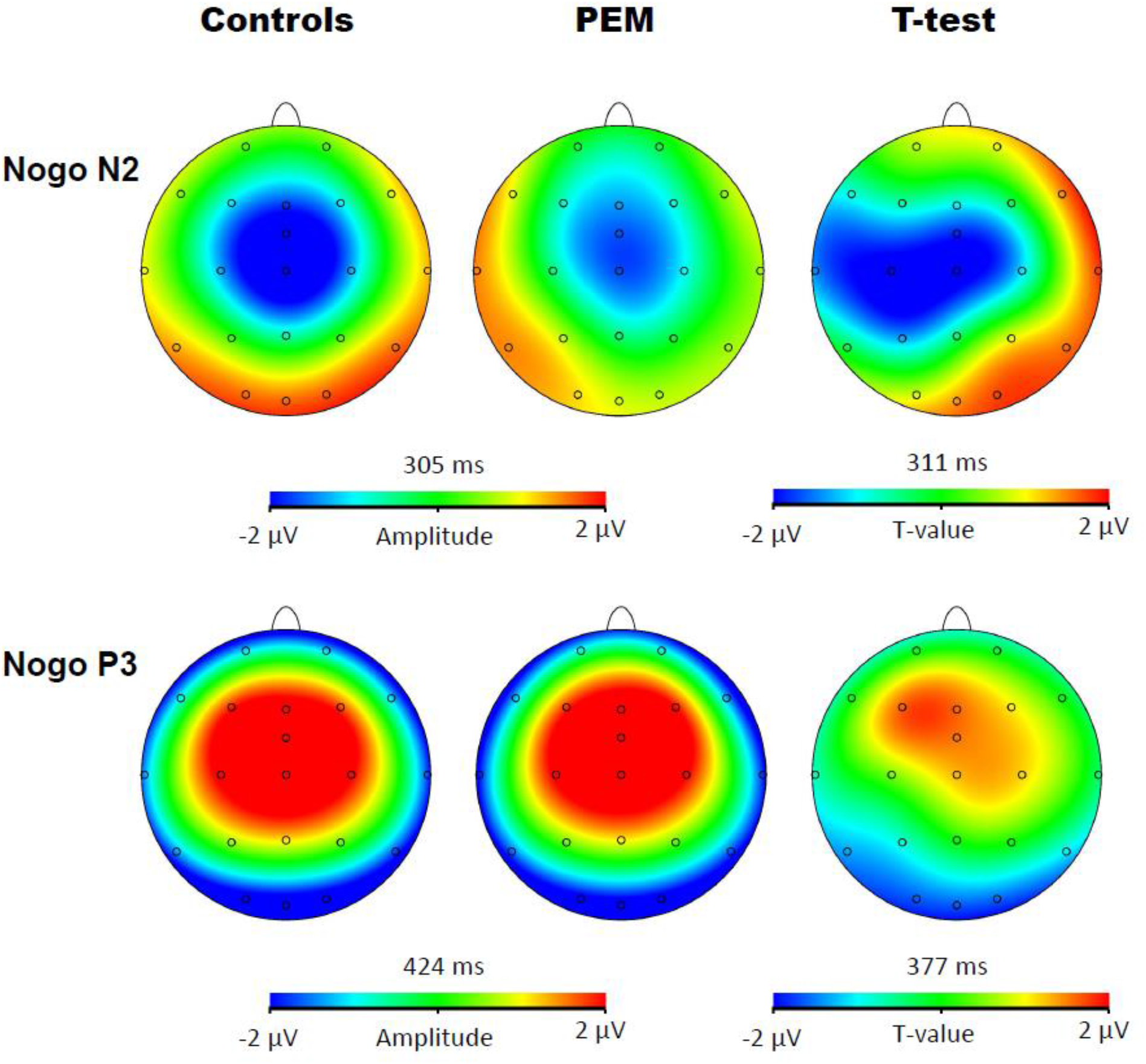
Topographic activation and topographic T-test (not corrected) of the difference of activation between PEM and control groups during No-Go condition

**Figure 3.**
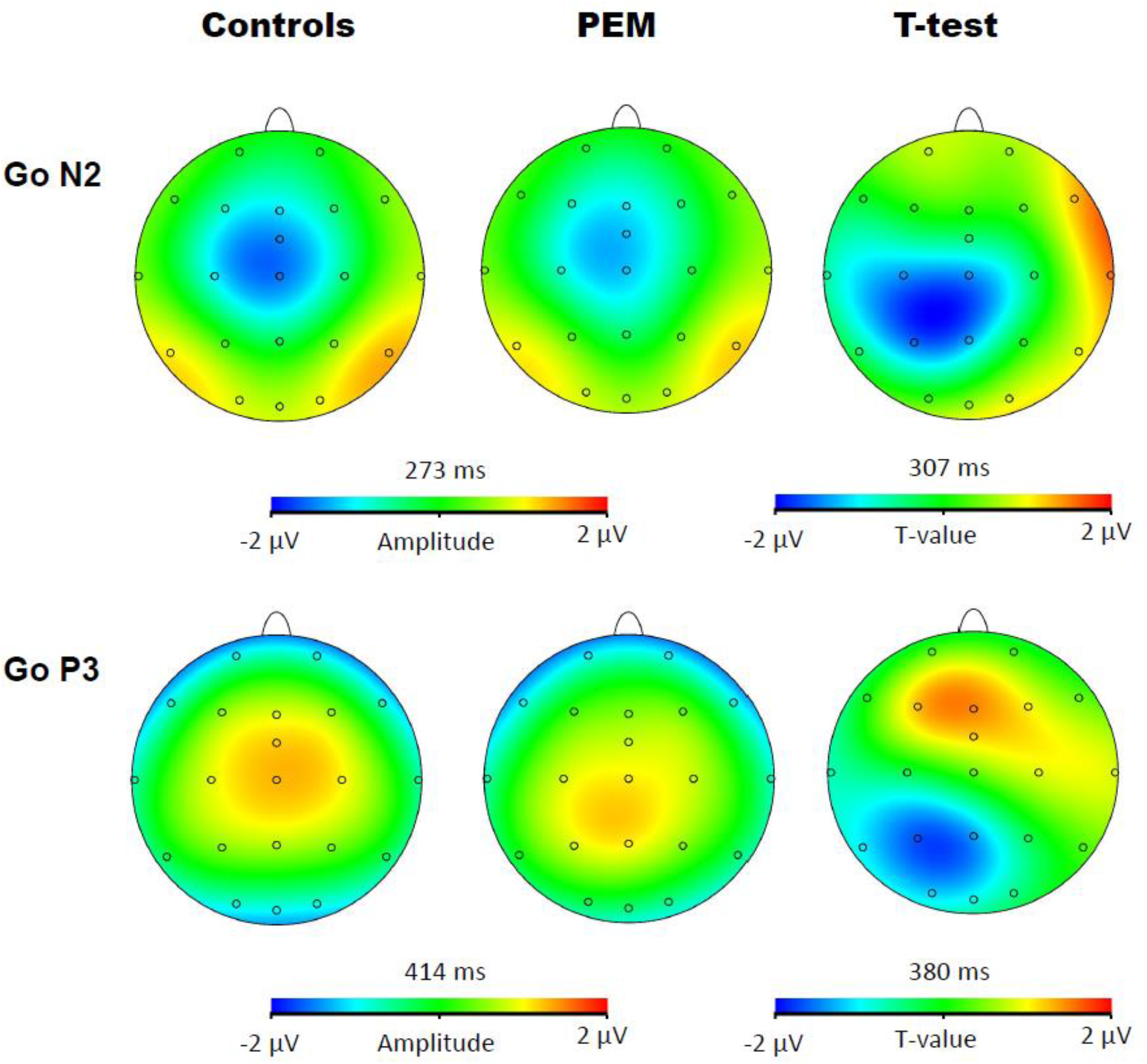
Topographic activation and topographic T-test (not corrected) of the difference of activation between PEM and control groups during Go condition

### 2.7. Statistical analyses

Data were analyzed SPSS statistical software, version 25. For the behavioral measures group differences in reaction time, Go accuracy and No-Go accuracy on the Go-No-Go task were analyzed using three separate independent sample t-tests with nutrition group as the between-subjects factor. For the ERP components, group differences were tested using four mixed ANOVAs [3 (Electrode: Fz, Cz, FCz) x 2 (Condition: No-Go, Go) x 2 (Nutrition Group: Controls, PEM)] performed separately for N2 and P3 and for their amplitude and latency. The significant p-value was set to *p*≤0.05. The extreme outliers, defined as a value that is over 3 times the interquartile range, were winsorized, as suggested by Wilcox and colleagues (2012). Greenhouse-Geisser adjustment was applied if the assumption of sphericity was violated (Mauchley’s test at *p*<0.05) and the Bonferroni correction was applied for multiple comparisons. A Mann-Whitney U-test was used if the Shapiro-Wilk normality test was violated. If nonparametric and parametric tests provided similar results, parametric tests were reported.

## 3. Results

### 3.1. Demographic characteristics

Demographic characteristics of the sample are reported in Table 1. The two nutrition groups did not differ in terms of age, gender or handedness, but there were significant differences in childhood standard of living with poor social status (higher HISP scores) in the PEM vs Control group.

**Table 1.** Demographic Characteristics of Participants - BNS Summer Study 2018

| Characteristic | PEM | Control | t-test / $\chi^2$ | *p* |
| --- | --- | --- | --- | --- |
| *N* | 24 | 29 |  |  |
| Males [*N* (%)] | 13 (54.17) | 14 (48.28) | 0,18 | 0,67 |
| Age (years) |  |  |  |  |
| Males (*SD*) | 49.2 (1.76) | 48.8 (1.99) | -0,58 | 0,57 |
| Females (*SD*) | 47.8 (1.74) | 48.5 (1.97) | 0,85 | 0,40 |
| Handedness [*N* left (%)] | 3 (12.50) | 3 (10.30) | 0,06 | 0,75 |
| Hollingshead Index of Social Position (ISP) |  |  |  |  |
| Education Index (*SD*) | 5.13 (0.34) | 3.79 (1.54) | -4,52 | < 0.0001 |
| Occupational Index (*SD*) | 6.25 (1.03) | 4.31 (1.58) | -5,36 | < 0.0001 |
| Total Index (*SD*) | 64.3 (7.55) | 45.3 (16.39) | -5,54 | < 0.0001 |
| ISP Category (*SD*) | 4.7 (0.48) | 3.4 (1.08) | -5,75 | < 0.0001 |

*Note.* Handedness statistical tests were done using Fisher Exact Test (non-parametric). There are 5 social position categories : (1) Upper, (2) Upper-middle, (3) Middle, (4) Lower-middle, and (5) Lower.

### 3.2. Behavioral results

Table 2 displays means and standard deviations of the behavioral measures. The nutrition groups differed on Go accuracy (*U*=462, *p*=0,041), with greater Go accuracy for the control group compared to the PEM group. However, there were no group differences in reaction time or No-Go accuracy.

**Table 2.** Mean and standard deviation of the behavioral measures for PEM and control group

| Measure | PEM ($N=24$) | | Control ($N=29$) | | Mann-Whitney $U$ | $p$ |
| --- | --- | --- | --- | --- | --- | --- |
| | $M$ | $SD$ | $M$ | $SD$ | | |
| Reaction time (ms) | 357,95 | 35,24 | 357,83 | 38,30 | 352,00 | 0,94 |
| Go accuracy (%) | 97,09 | 3,22 | 98,25 | 2,59 | 462,00 | 0,04 |
| No-Go accuracy (%) | 76,64 | 12,11 | 80,21 | 12,87 | 420,00 | 0,20 |

### 3.3. ERP Results: N2

Figure 4 and 5 show the waveform of the N2 and P3 components for each nutrition group during the No-Go and Go conditions, respectively.

**Figure 4.**
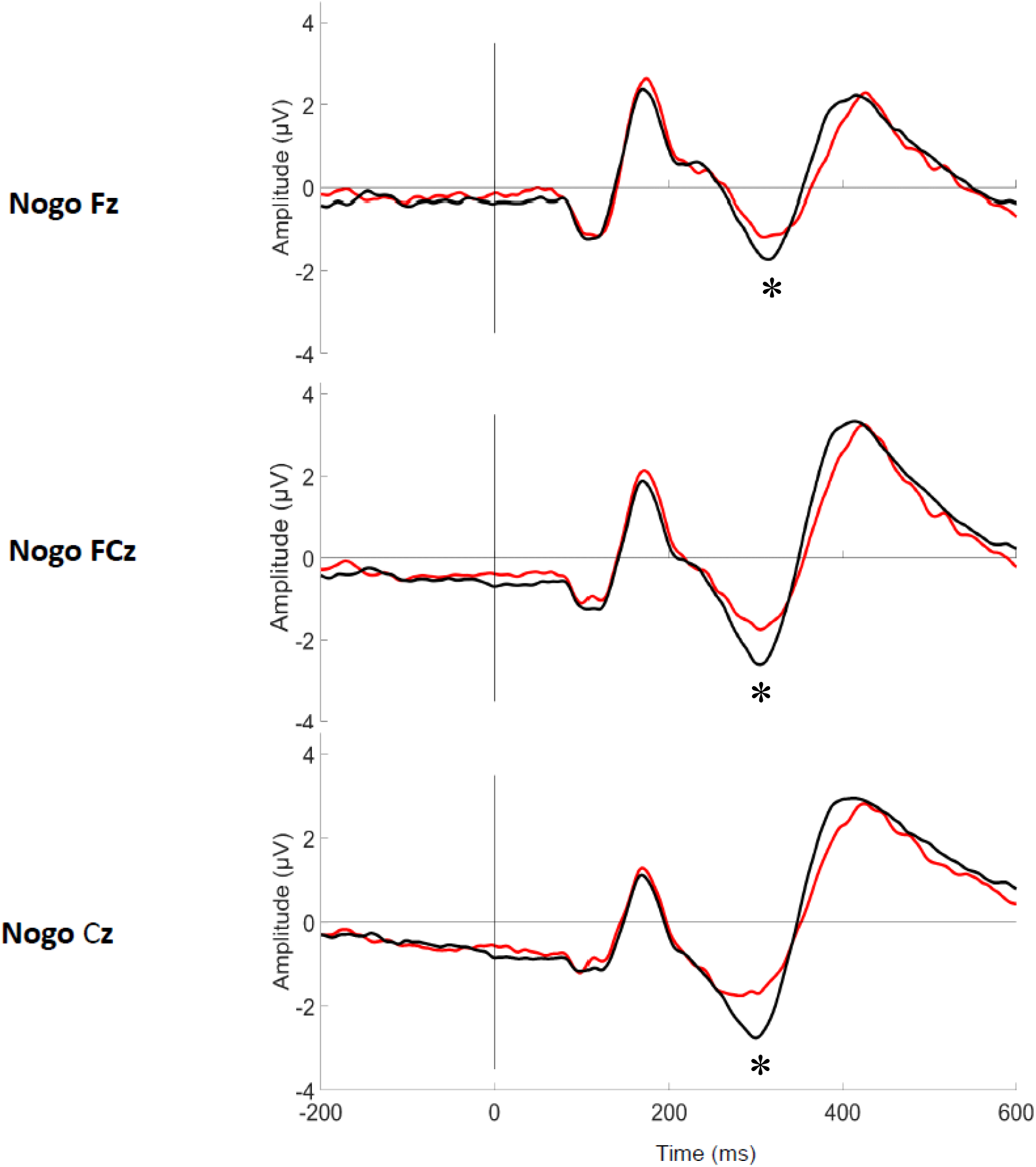
Grand average waveform of the N2 and P3 components during No-Go condition for each group

**Figure 5.**
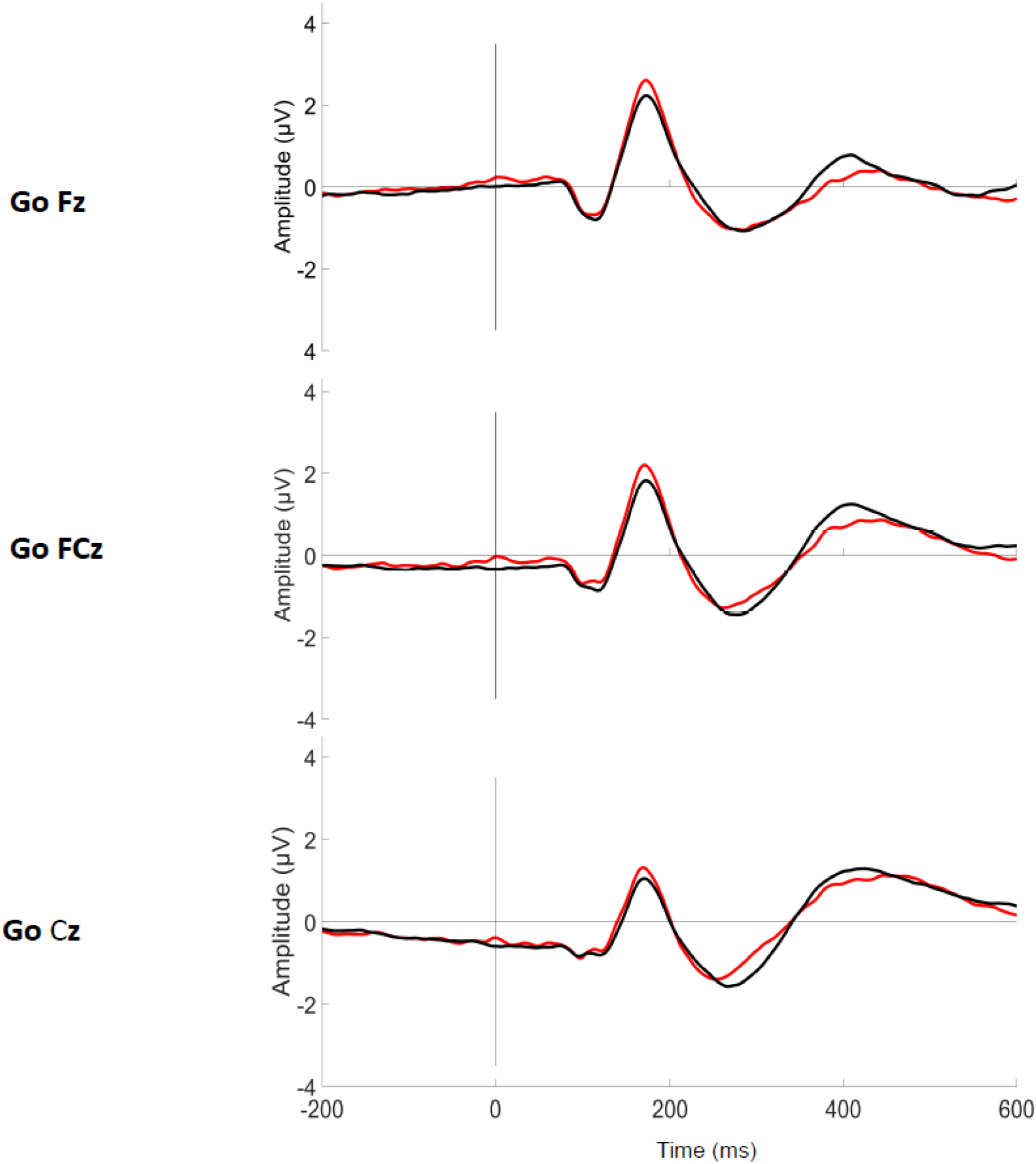
Grand average waveform of the N2 and P3 components during Go condition for each group

#### 3.3.1. Amplitude

A three-way ANOVA revealed a main effect of Electrodes (*F* (2,102) = 7.745, *p*=0.004, η_p_^2^=0.132) with pairwise comparisons indicating a smaller amplitude for Fz when compared with FCz (*p*<0.001) and Cz, although the latter difference did not achieve statistical significance (*p*=0.060). There was also a main effect of Condition (*F* (1,51) = 30.219, *p*<0.001, η_p_^2^=0.372) with the No-Go condition generating a larger N2 amplitude than the Go condition, according to pairwise comparisons. Significant interactions between Condition and Nutrition Group (*F* (1,51) = 5.404, *p*=0.024, η_p_^2^=0.096) and between Electrode and Condition (*F* (2,102) = 5.061, *p*=0.020, η_p_^2^=0.090) were found. Appropriate t-tests and repeated measures ANOVA were conducted to further explore these interactions.

For the Condition x Nutrition Group interaction, independent t-tests revealed a significant difference of the N2 amplitude between the two groups but only in the No-Go condition [No-Go: *t*(51) = 1.959, *p*=0.056; Go: (*t*(51) = 0.147, *p*=0.883]. During this condition, the amplitude of N2 was significantly larger for the control group compared to the PEM group.

For the Electrode x Condition interaction, the repeated measure ANOVA revealed a smaller amplitude for Fz compared to FCz (*p*<0,001) and Cz (*p*=0,035) during the No-Go condition (*F*(2,104)=9,103, *p*=0,002), but only a smaller amplitude for Fz compared to FCz (*p*=0,027) during the Go condition (*F*(2,104)=3,479, *p*=0,058).

#### 3.3.2. Latency

A three-way ANOVA showed a main effect of Electrodes (*F* (2,102) = 13.943, *p*<0.001, η_p_^2^=0.215) with pairwise comparisons indicating a shorter latency for Cz compared to FCz (*p*=0.001) and Fz (*p*<0.001). There was also a main effect of Condition (*F* (1,51) = 22.524, *p*<0.001, η_p_^2^=0.306) with the No-Go condition generating a longer N2 latency than the Go condition. No interaction was significant, and there was no significant nutrition group effect.

### 3.4. ERP Results: P3

#### 3.4.1. Amplitude

The three-way ANOVA revealed a main effect of Electrodes (*F* (2,102) = 25.811, *p*<0.001, η_p_^2^=0.336) with pairwise comparisons indicating a smaller amplitude for Fz compared to FCz (*p*<0.001) and Cz (*p*=0.001). There was also a main effect of Condition (*F* (1,51) = 195.127, *p*<0.001, η_p_^2^=0.793) with the No-Go condition generating a larger P3 amplitude than the Go condition, according to pairwise comparisons. There was also a significant interaction between Electrode and Condition (*F* (2,102) = 7.667, *p*=0.004, η_p_^2^=0.131). A repeated measures ANOVA was conducted as posthoc analysis to further explore this interaction. For the No-Go condition (*F*(2,104)=18.912, *p*<0.001, η_p_^2^=0.267), P3 amplitude was significantly larger at FCz compared to Fz (*p*<0.001) and Cz (*p*=0.014), and was larger at Cz compared to Fz (*p*=0.027). For the Go condition (*F*(2,104)=26.076, *p*<0.001, η_p_^2^=0.334), P3 amplitude was smaller at Fz compared to FCz (*p*<0.001) and Cz (*p*<0.001). There was no significant effect of Nutrition Group on the measure.

#### 3.4.2. Latency

No significant main effect or interaction was obtained for the P3 latency. Table 3, 4 and 5 display means and standard deviations of the ERP measures for each condition for Fz, Cz and FCz, respectively.

**Table 3.**
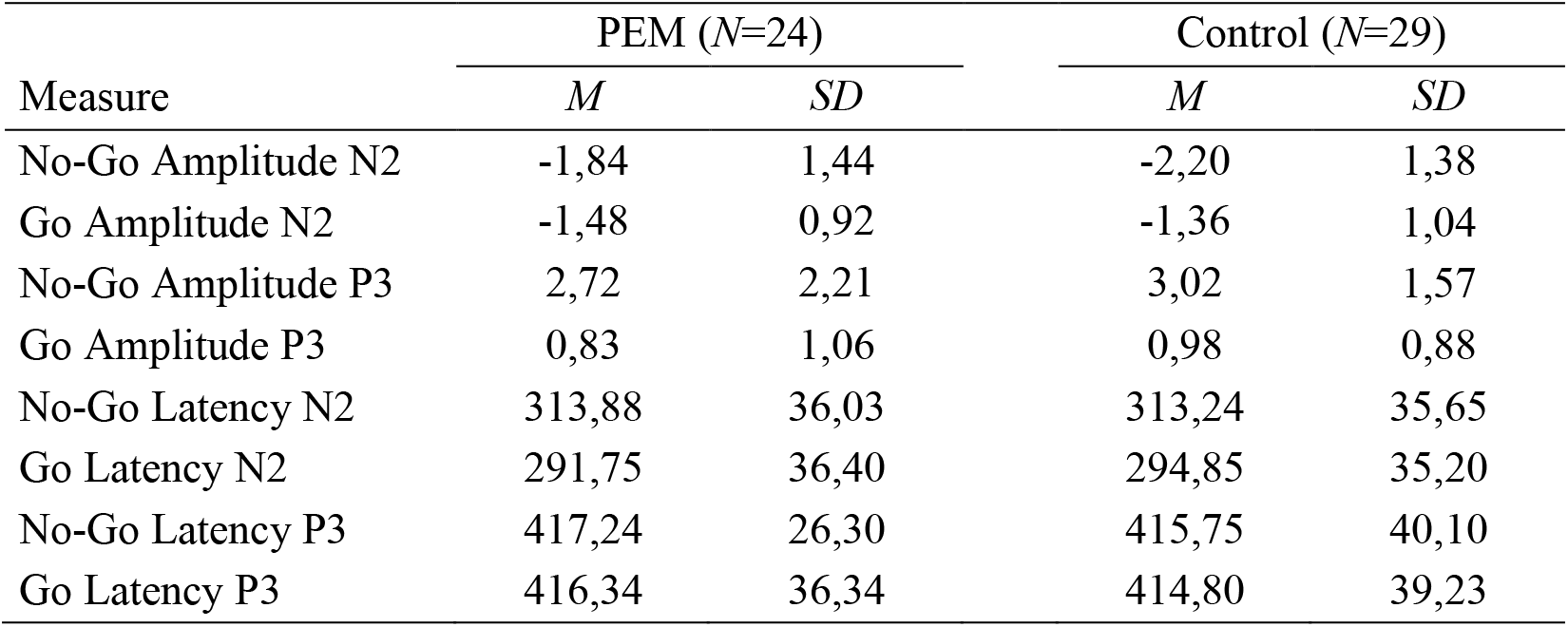
Mean and standard deviation of the ERP measures for Fz electrode

| Measure | PEM (N=24) |  | Control (N=29) |  |
| --- | --- | --- | --- | --- |
|  | *M* | *SD* | *M* | *SD* |
| No-Go Amplitude N2 | -1,84 | 1,44 | -2,20 | 1,38 |
| Go Amplitude N2 | -1,48 | 0,92 | -1,36 | 1,04 |
| No-Go Amplitude P3 | 2,72 | 2,21 | 3,02 | 1,57 |
| Go Amplitude P3 | 0,83 | 1,06 | 0,98 | 0,88 |
| No-Go Latency N2 | 313,88 | 36,03 | 313,24 | 35,65 |
| Go Latency N2 | 291,75 | 36,40 | 294,85 | 35,20 |
| No-Go Latency P3 | 417,24 | 26,30 | 415,75 | 40,10 |
| Go Latency P3 | 416,34 | 36,34 | 414,80 | 39,23 |

**Table 4.** Mean and standard deviation of the ERP measures for FCz electrode

| Measure | PEM (N=24) |  | Control (N=29) |  |
| --- | --- | --- | --- | --- |
|  | *M* | *SD* | *M* | *SD* |
| No-Go Amplitude N2 | -2,04 | 1,78 | -3,07 | 1,53 |
| Go Amplitude N2 | -1,53 | 0,97 | -1,72 | 1,23 |
| No-Go Amplitude P3 | 3,38 | 1,52 | 3,70 | 1,80 |
| Go Amplitude P3 | 1,33 | 0,97 | 1,53 | 0,97 |
| No-Go Latency N2 | 305,09 | 37,43 | 301,19 | 30,39 |
| Go Latency N2 | 279,70 | 41,46 | 281,92 | 36,56 |
| No-Go Latency P3 | 417,48 | 24,47 | 420,39 | 44,71 |
| Go Latency P3 | 416,10 | 37,53 | 413,46 | 40,61 |

**Table 5.** Mean and standard deviation of the ERP measures for Cz electrode

| Measure | PEM (N=24) |  | Control (N=29) |  |
| --- | --- | --- | --- | --- |
|  | *M* | *SD* | *M* | *SD* |
| No-Go Amplitude N2 | -2,19 | 1,63 | -3,06 | 1,52 |
| Go Amplitude N2 | -1,61 | 1,02 | -1,66 | 1,23 |
| No-Go Amplitude P3 | 3,74 | 2,19 | 4,24 | 1,76 |
| Go Amplitude P3 | 1,21 | 0,91 | 1,55 | 1,02 |
| No-Go Latency N2 | 307,94 | 36,42 | 310,01 | 23,93 |
| Go Latency N2 | 287,60 | 37,51 | 291,49 | 33,66 |
| No-Go Latency P3 | 426,76 | 34,67 | 416,15 | 39,51 |
| Go Latency P3 | 415,93 | 37,06 | 412,85 | 38,25 |

## 4. Discussion

The main objective of this study was to compare the brain activity of adults who experienced moderate to severe PEM during the first year of life and healthy controls with no histories of malnutrition using a Go-No-Go inhibition task. We hypothesized that the PEM group would show altered neural response associated with attention and inhibition (N2 and P3 components) during the task. The hypothesis was partially confirmed, since we observed a reduction of N2 amplitude during the No-Go condition in the PEM group compared to the control group, but no difference in P3. Overall, results of the Go-No-Go task revealed a main effect of Condition for both components (N2 and P3) in both groups (PEM and Control). This is typical result for a Go-No-Go task, indicating that the No-Go condition induces a genuine inhibition response process which is delaying the onset and amplifying the N2 and P3 components (Bokura et al., 2001; Eimer, 1993; Schmüser et al., 2016).

The N2 amplitude was smaller in the PEM group compared to the control group during the No-Go condition. This result has also been found in adults and children with ADHD (Brandeis et al., 2002; Fallgatter et al., 2004; Gow et al., 2012; Woltering et al., 2013). There is debate amongst the scientific community on the cognitive process underlying the N2 component. Although some authors argue that N2 is related to the inhibition process (i.e. cancellation of a planned or prepotent response; Eimer, 1993; Falkenstein et al., 1999), more recent studies point to an association between N2 and conflict monitoring (i.e. conflict between the prepotent response and required response; Donkers and van Boxtel, 2004; Enriquez-Geppert et al., 2010; Groom and Cragg, 2015; Huster et al., 2013). According to the conflict monitoring hypothesis, an inhibition task such as Go-No-Go should evoke a N2 component because of the unbalanced ratio between Go and No-Go trials, which would lead to the creation of a prepotent response (Go) that conflicts with the infrequent inhibition of this response (No-Go) and not because of the inhibition process *per se* (Braver et al., 2001). Conflict monitoring is however closely related to attention since it is responsible for triggering cognitive control changes by adjusting attention levels to optimize performance and prevent subsequent conflict (Botvinick et al., 2001). According to these models, our results can be interpreted as an impairment in conflict monitoring and/or attention following early childhood malnutrition.

Surprisingly, the ERP results showed no difference between the two nutrition groups in P3 amplitude or latency. P3 component is usually considered to be a marker of response inhibition processing and evaluation (Groom and Cragg, 2015; Huster et al., 2013). In studies of ADHD using a Go-no-Go task, both components, N2 and P3, are typically altered (Johnstone et al., 2013; Woltering et al., 2013). However, the reduced N2 amplitude in the PEM group and similar P3 amplitude between groups is in line with the behavioral results. Indeed, in the current study, Go accuracy was significantly lower in the PEM group compared to the control group, revealing that adults who suffered from early childhood malnutrition committed more omission errors than controls. No difference in commission error rate was found. Omission errors are usually attributed to impairments in attention and vigilance whereas commission errors are associated with inhibition deficits. Therefore, early childhood malnutrition may be associated with diminished attention and vigilance rather than altered inhibition skills. Furthermore, no differences in reaction times were found between our groups. Slower reaction times have been consistently reported in populations with attentional and inhibition difficulties and are associated with hyperactivity and impulsivity (Barkley et al., 2008; McLoughlin et al., 2010; Rubia et al., 2007; Schmüser et al., 2016; Tamm et al., 2012; Wiersema et al., 2006). Overall, the behavioral results suggest attention deficits with normal inhibition skills in adults who experienced early childhood malnutrition. Nevertheless, we cannot rule out that the PEM group developed compensatory mechanisms to overcome inhibition difficulties.

The ERP and behavioral results are in line with the persistent attention deficits previously reported in our cohort during childhood, adolescence and adulthood (Galler et al., 2012c, 1990, 1983a; Galler and Ramsey, 1989; Peter et al., 2016). Attention deficits have also been reported in other studies assessing the neurocognitive profile of malnourished populations (Kar et al., 2008; Kesari et al., 2010; Richardson et al., 1972; Wang et al., 2016). Although the PEM group had persistent attention deficits in the prior BNS publications (e.g. 69% had at least one score falling within the clinical range for attention disorder at the Continuous Performance Test or CPT), only 8% obtained a clinical diagnosis of ADHD (Galler et al., 2012c). Attention deficits seem highly more prevalent in our cohort than hyperactive/impulsive ADHD symptoms. Interestingly, a recent study assessing neuropsychological functions in the BNS cohort showed that cognitive flexibility was more altered than inhibition (Waber et al., 2014a). This neuropsychological profile could explain why we did not find any electrophysiological marker of inhibition deficits (P3 or commission errors).

The altered electrophysiological marker found in this study is also coherent with literature on brain function effects following childhood malnutrition. Indeed, several studies show that childhood malnutrition is associated with slowing of the EEG’s dominant rhythm in infancy (Baraitser and Evans, 1969; Gladstone et al., 2014; Griesel et al., 1990; Stoch et al., 1963) and electrophysiological alterations that persist in childhood even with food rehabilitation (Bartel et al., 1979; Taboada-Crispi et al., 2018). Barnet and colleagues (1978; 2007) also found ERP abnormalities following childhood malnutrition with a higher relative abnormality of their auditory evoked potentials compared to controls. We expand knowledge in this field by showing brain function deficits still perceptible in adulthood.

This study has several limitations. First, the small sample size does not allow us to conclude with certainty that the results were not due to lack of statistical power. Nevertheless, the effect size is moderate for the N2 amplitude difference between the groups (*d*=0.55). Also, due to the small sample size, we could not adjust our statistical analyses for age, gender and handedness. Even though we found no differences between the two groups on those variables, we cannot exclude that these variables might have had an effect on our results. Additionally, the results reported here are specific to the N2 and P3 electrophysiological markers. Potential other electrophysiological differences between our groups might exist and would allow us to identify additional brain markers of early childhood malnutrition (e.g. time frequency, source level, connectivity). Indeed, using source localization analysis on our data would allow to explore more precisely at the source level the brain function temporal dynamics of the executive function. However, at least 64 electrodes would be needed to apply source analysis. Further analyses will be performed to better characterize and compare brain function in both groups.

## 5. Conclusion

In sum, this study shows that adults who experienced early childhood malnutrition in the first year of life demonstrate different brain response patterns during a response inhibition task compared to healthy controls. This adds to the existent literature on cognitive and neural outcomes following early childhood malnutrition, suggesting that attention and conflict monitoring, two cognitive control processes, are still altered in adulthood. Malnutrition can have deleterious effects on cognition, physical and mental health, behavior and brain function even if restricted to the first year of life. Considering the impact of persisting cognitive alterations on the quality of life, more research is needed to better characterize the brain markers and clinical profile associated with early childhood malnutrition in order to develop a disease progression model applicable to various vulnerable populations.

## Statement of Ethical Standards

This study has been performed in accordance with the ethical standards laid down in the 1964 Declaration of Helsinki and its later amendments. All study participants provided written informed consent and were compensated for their time and travel expenses. This study was approved by the Massachusetts General Hospital IRB (IRB Protocol 2015P000329/MGH), Hôpital Sainte-Justine and Centro de Neurosciencias de Cuba’s (CNEURO) Ethics’ committees.

## Declaration of Interest

The authors declare no competing interests.

## Funding

This research was conducted in cooperation with the Ministry of Health of Barbados and was supported by grants from the QBIN (AG), the National Institutes of Health (JRG; HD060986) and also by grants from the National Nature Science Foundation of China (PVS, MB; Nos. 61673090 and 81330032) and the Nestle Foundation (PVS).

## Data and Code Availability Statement

The data that support the findings of this study are available from the corresponding author, [AG], upon reasonable request.

## Notes

### Competing Interest Statement

The authors have declared no competing interest.

